## Supplemental Tables and Figures for "Optimization of systemic AAV9 gene therapy in Niemann-Pick disease, type C1 mice"

#### Table S1

| Antibody | Company | Catalog Number | Dilution | Method |
|---|---|---|---|---|
| **α-β-actin (mouse IgG)** | ThermoFisher Invitrogen | 15G5A11/E2 | 1:10000 | western blot |
| **α-Calbindin (mouse IgG)** | Sigma | C9848 | 1:750<br>1:1000 | IF<br>western blot |
| **α-Calbindin (rabbit IgG)** | Abcam | ab229915 | 1:750 (fixed, free-floating sections and FFPE sections) | IF |
| **α-CD68 (rat IgG2a)** | BioRad | MCA1957 | 1:500 | IF |
| **α-CD68 (rabbit IgG)** | Abcam | ab125212 | 1:1000 | IHC, IF, western blot |
| **α-GFAP (mouse IgG)** | Sigma | G3893 | 1:1000 (fixed, free-floating sections and FFPE sections)<br>1:2000 | IF<br>western blot |
| **α-IBA1 (rabbit IgG)** | Wako Chemicals | 019-19741 | 1:500 (fixed, free-floating sections) or 1:750 (FFPE) | IF |
| **α-NPC1 (monoclonal rabbit IgG)** | Abcam | ab134113 | 1:2000 | western blot |
| **α-β-III-Tubulin** | R&D Systems | MAB1195 | 1:3000 | western blot |
| **Donkey anti-mouse IgG IRDye 680RD** | LI-CORbio | 926-68072 | 1:20000 | western blot |
| **Donkey anti-mouse IgG IRDye 800CW** | LI-CORbio | 926-32213 | 1:20000 | western blot |
| **Goat anti-rabbit IgG Biotinylated** | Vector Laboratories | BA-1000 | 1:300 | IHC |
| **Goat anti-mouse IgG AlexaFluor 488 or 594** | ThermoFisher Invitrogen | A11029 (488) or A11005 (594) | 1:350 | IF |
| **Goat anti-rabbit IgG AlexaFluor 488 or 594** | ThermoFisher Invitrogen | A11034 (488) or A11037 (594) | 1:350 | IF |
| **Goat anti-rat IgG Alexa Fluor 488 or 594** | ThermoFisher Invitrogen | A11006 (488) or A11007 (594) | 1:350 | IF |

**Table S2**

**A**

| **Treatment** | **Sample Size (n)** | **Median Survival (weeks)** | **Significance (Log-rank test)** |
|---|---|---|---|
| *Npc1*$^{m1N}$ Saline | 15 | 10.6 | |
| *Npc1*$^{m1N}$ Low 7.87x10$^{12}$ vg/kg | 10 | 11.4 | vs. Saline, ***P=0.0049*** |
| *Npc1*$^{m1N}$ Medium 1.28x10$^{14}$ vg/kg | 24 | 21.5 | vs. Saline, ***P<0.0001***<br>vs. Low, ***P<0.0001*** |
| *Npc1*$^{m1N}$ High 3.06x10$^{14}$ vg/kg | 8 | 34.6 | vs. Saline, ***P<0.0001***<br>vs. Low, ***P<0.0001***<br>vs. Medium, P=0.0266 |

**B**

| **Treatment** | **Sample Size (n)** | **Median Survival (weeks)** | **Significance (Log-rank test)** |
|---|---|---|---|
| *Npc1*$^{m1N}$ Saline | 15 | 10.6 | |
| *Npc1*$^{m1N}$ Med AAV9 at **4 weeks old** | 24 | 21.5 | vs. Saline, ***P<0.0001*** |
| *Npc1*$^{m1N}$ Med AAV9 at **6 weeks old** | 20 | 13.2 | vs. Saline, ***P<0.0001***<br>vs. 4 weeks old, ***P=0.0030*** |
| *Npc1*$^{m1N}$ Med AAV9 at **8 weeks old** | 20 | 11.9 | vs. Saline, ***P=0.0006***<br>vs. 4 weeks old, ***P<0.0001***<br>vs. 6 weeks old, ***P=0.0003*** |

#### Table S3

**A**

| Treatment | Sample Size (n) | Significance (Tukey's Multiple Comparisons Test, 6-9 weeks) | Significance (Tukey's Multiple Comparisons Test, 9-12 weeks) |
|---|---|---|---|
| *Npc1[m1N]* Saline | 14 | | |
| *Npc1[m1N]* Low<br>7.87x10[12] vg/kg | 10 | vs. Saline, P=0.9466 | vs. Saline, P=0.9976 |
| *Npc1[m1N]* Medium<br>1.28x10[14] vg/kg | 13 | vs. Saline, P=0.0196<br>vs. Low, ***P=0.0069*** | vs. Saline, ***P<0.0001***<br>vs. Low, ***P<0.0001*** |
| *Npc1[m1N]* High<br>3.06x10[14] vg/kg | 8 | vs. Saline, ***P<0.0001***<br>vs. Low, ***P<0.0001***<br>vs. Medium, P=0.0129 | vs. Saline, ***P<0.0001***<br>vs. Low, ***P<0.0001***<br>vs. Medium, **P=0.0009** |
| *Npc1[++]* | 21 | vs. Saline, ***P<0.0001***<br>vs. Low, ***P<0.0001***<br>vs. Medium, ***P=0.0003***<br>vs. High, P=0.9104 | vs. Saline, ***P<0.0001***<br>vs. Low, ***P<0.0001***<br>vs. Medium, ***P<0.0001***<br>vs. High, P=0.5073 |

**B**

| Treatment | Sample Size (n) | Significance (Tukey's Multiple Comparisons Test, 6-9 weeks) | Significance (Tukey's Multiple Comparisons Test, 9-12 weeks) |
|---|---|---|---|
| *Npc1[m1N]* Saline | 14 | | |
| *Npc1[m1N]* Med AAV9 at **4 weeks old** | 13 | vs. Saline, ***P<0.0001*** | vs. Saline, ***P<0.0001*** |
| *Npc1[m1N]* Med AAV9 at **6 weeks old** | 20 | vs. Saline, P=0.7788<br>vs. 4 weeks old, ***P=0.0010*** | vs. Saline, P=0.1909<br>vs. 4 weeks old, ***P=0.0002*** |
| *Npc1[m1N]* Med AAV9 at **8 weeks old** | 20 | vs. Saline, P=.9984<br>vs. 4 weeks old, ***P<0.0001***<br>vs. 6 weeks old, P=0.5116 | vs. Saline, P>0.9999<br>vs. 4 weeks old, ***P<0.0001***<br>vs. 6 weeks old, P=0.0747 |
| *Npc1[++]* | 21 | vs. Saline, ***P<0.0001***<br>vs. 4 weeks old, P=0.2389<br>vs. 6 weeks old, ***P<0.0001***<br>vs. 8 weeks old, ***P<0.0001*** | vs. Saline, ***P<0.0001***<br>vs. 4 weeks old, ***P<0.0001***<br>vs. 6 weeks old, ***P<0.0001***<br>vs. 8 weeks old, ***P<0.0001*** |

**Table S4**

**A**

| AAV9 Dose Study (Sample size for weight curves) | | | | | |
|---|---|---|---|---|---|
| Treatment | *Npc1*$^{m1N}$ saline | *Npc1*$^{m1N}$ low | *Npc1*$^{m1N}$ med | *Npc1*$^{m1N}$ high | *Npc1*$^{+/+}$ untreated |
| Male | 8 | 6 | 10 | 4 | 11 |
| Female | 7 | 4 | 14 | 4 | 10 |
| **Total** | **15** | **10** | **24** | **8** | **21** |

**B**

| AAV9 Age at Injection Study (Sample size for weight curves) | | | | | |
|---|---|---|---|---|---|
| Treatment | *Npc1*$^{m1N}$ saline | *Npc1*$^{m1N}$ med 4 wks | *Npc1*$^{m1N}$ med 6 wks | *Npc1*$^{m1N}$ med 8 wks | *Npc1*$^{+/+}$ untreated |
| Male | 8 | 10 | 11 | 12 | 12 |
| Female | 7 | 14 | 9 | 8 | 10 |
| **Total** | **15** | **24** | **20** | **20** | **22** |

**C**

| AAV9 Hypomorphic (I1061T) Model Study (Sample size for weight curves) | | | |
|---|---|---|---|
| Treatment | *Npc1*$^{I1061T}$ saline | *Npc1*$^{I1061T}$ med | *Npc1*$^{+/+}$ untreated |
| Male | 5 | 7 | 6 |
| Female | 6 | 8 | 9 |
| **Total** | **11** | **15** | **15** |

Figure S1

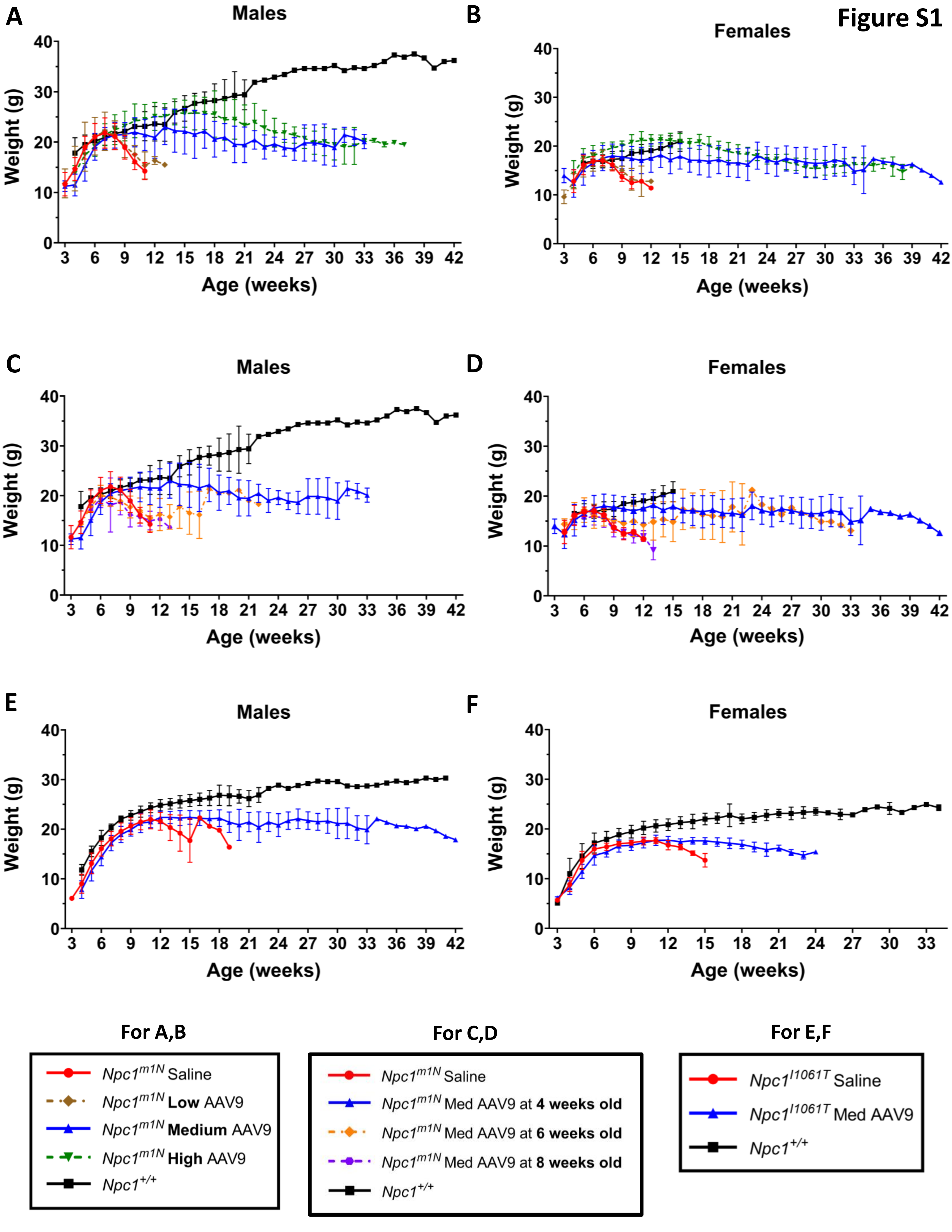
A
Males
B
Females
C
Males
D
Females
E
Males
F
Females
Weight (g)
Age (weeks)
For A,B
Npc1^m1N Saline
Npc1^m1N Low AAV9
Npc1^m1N Medium AAV9
Npc1^m1N High AAV9
Npc1^+/+
For C,D
Npc1^m1N Saline
Npc1^m1N Med AAV9 at 4 weeks old
Npc1^m1N Med AAV9 at 6 weeks old
Npc1^m1N Med AAV9 at 8 weeks old
Npc1^+/+
For E,F
Npc1^I1061T Saline
Npc1^I1061T Med AAV9
Npc1^+/+

**Figure S2**

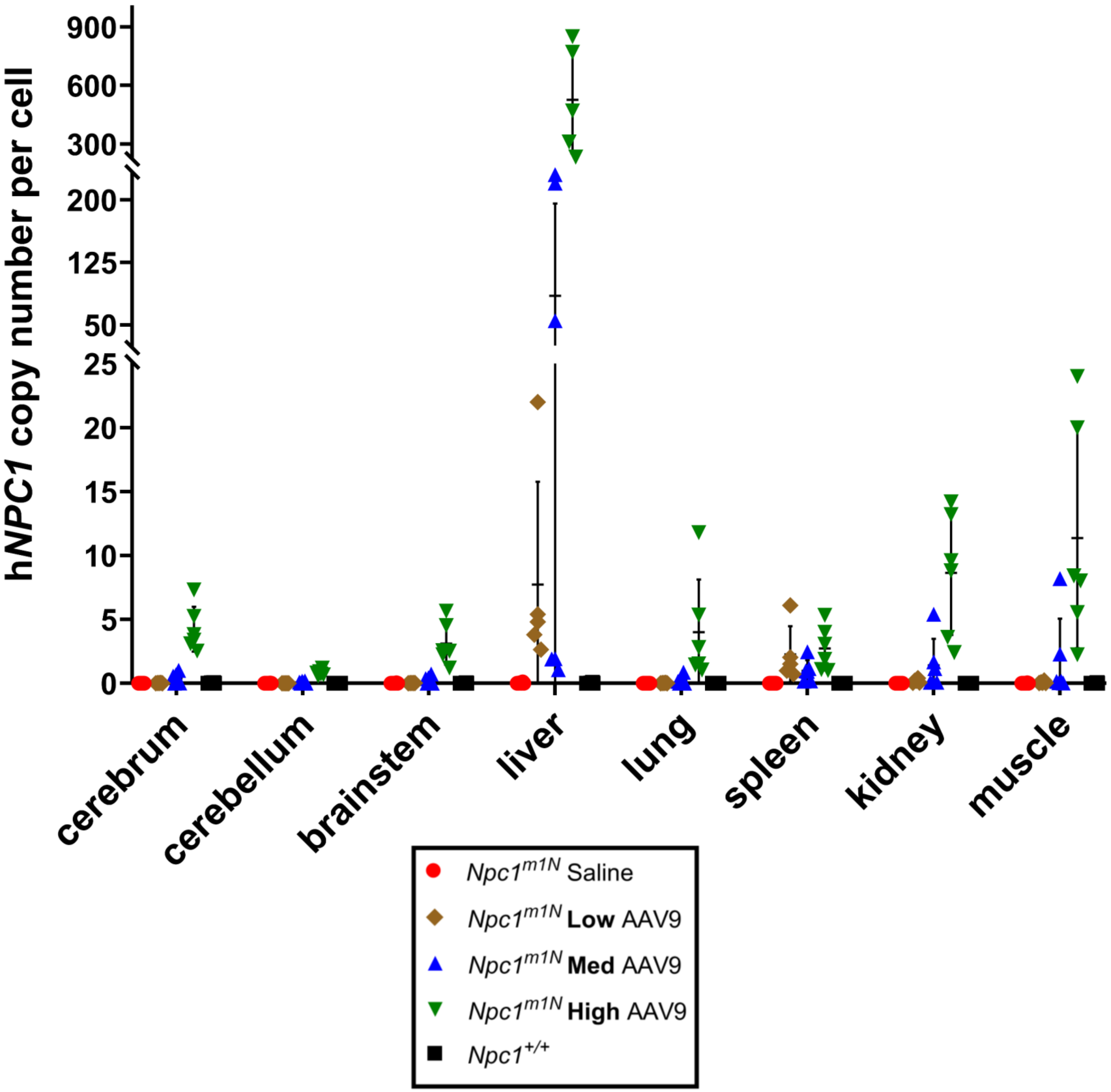

hNPC1 copy number per cell
900
600
300
200
125
50
25
20
15
10
5
0
cerebrum
cerebellum
brainstem
liver
lung
spleen
kidney
muscle
Npc1m1N Saline
Npc1m1N Low AAV9
Npc1m1N Med AAV9
Npc1m1N High AAV9
Npc1+/+

**Figure S3**

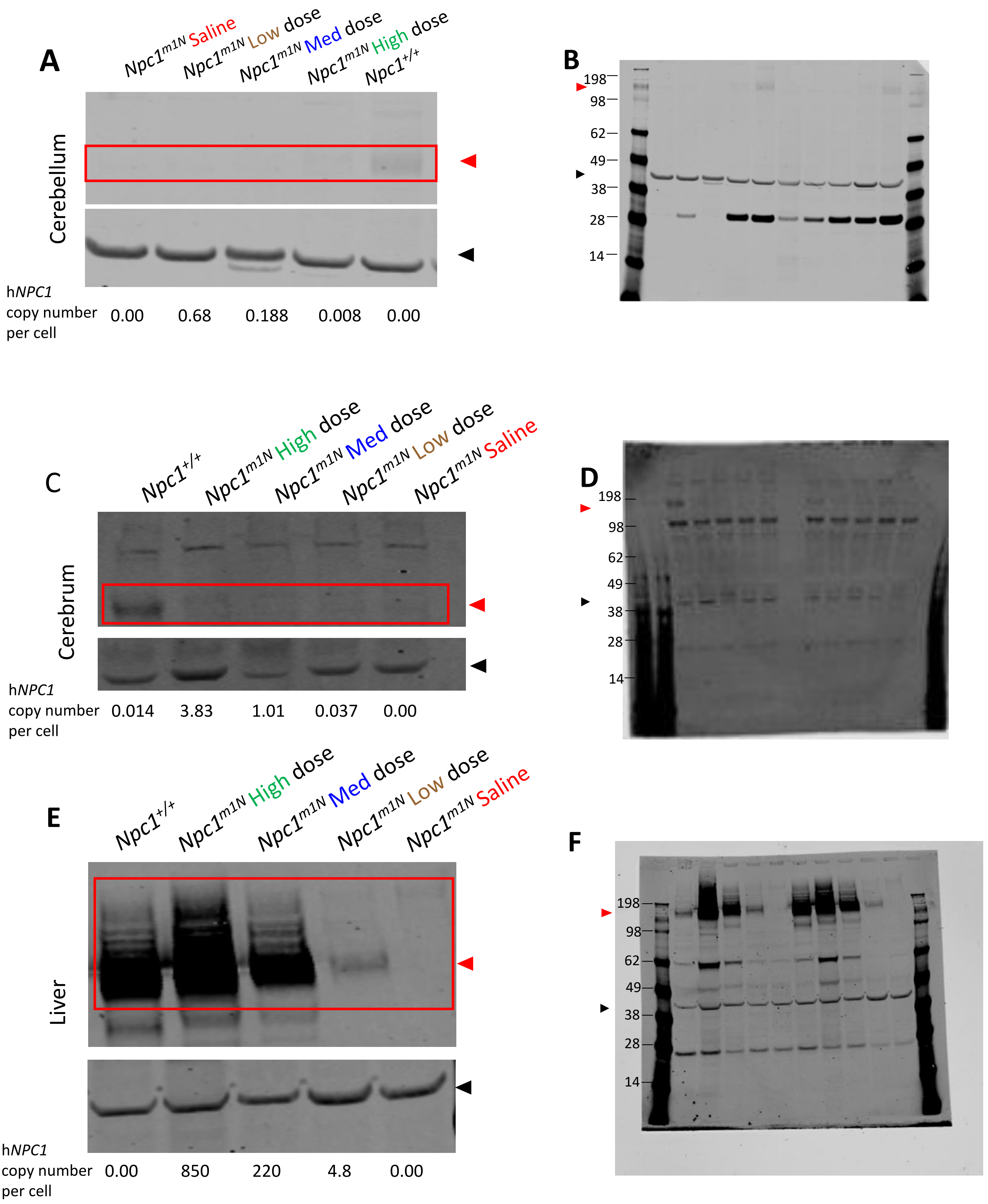
A
Npc1m1N Saline
Npc1m1N Low dose
Npc1m1N Med dose
Npc1m1N High dose
Npc1+/+
Cerebellum
hNPC1 copy number per cell
0.00
0.68
0.188
0.008
0.00
B
198
98
62
49
38
28
14
C
Npc1+/+
Npc1m1N High dose
Npc1m1N Med dose
Npc1m1N Low dose
Npc1m1N Saline
Cerebrum
hNPC1 copy number per cell
0.014
3.83
1.01
0.037
0.00
D
198
98
62
49
38
28
14
E
Npc1+/+
Npc1m1N High dose
Npc1m1N Med dose
Npc1m1N Low dose
Npc1m1N Saline
Liver
hNPC1 copy number per cell
0.00
850
220
4.8
0.00
F
198
98
62
49
38
28
14

### Figure S4

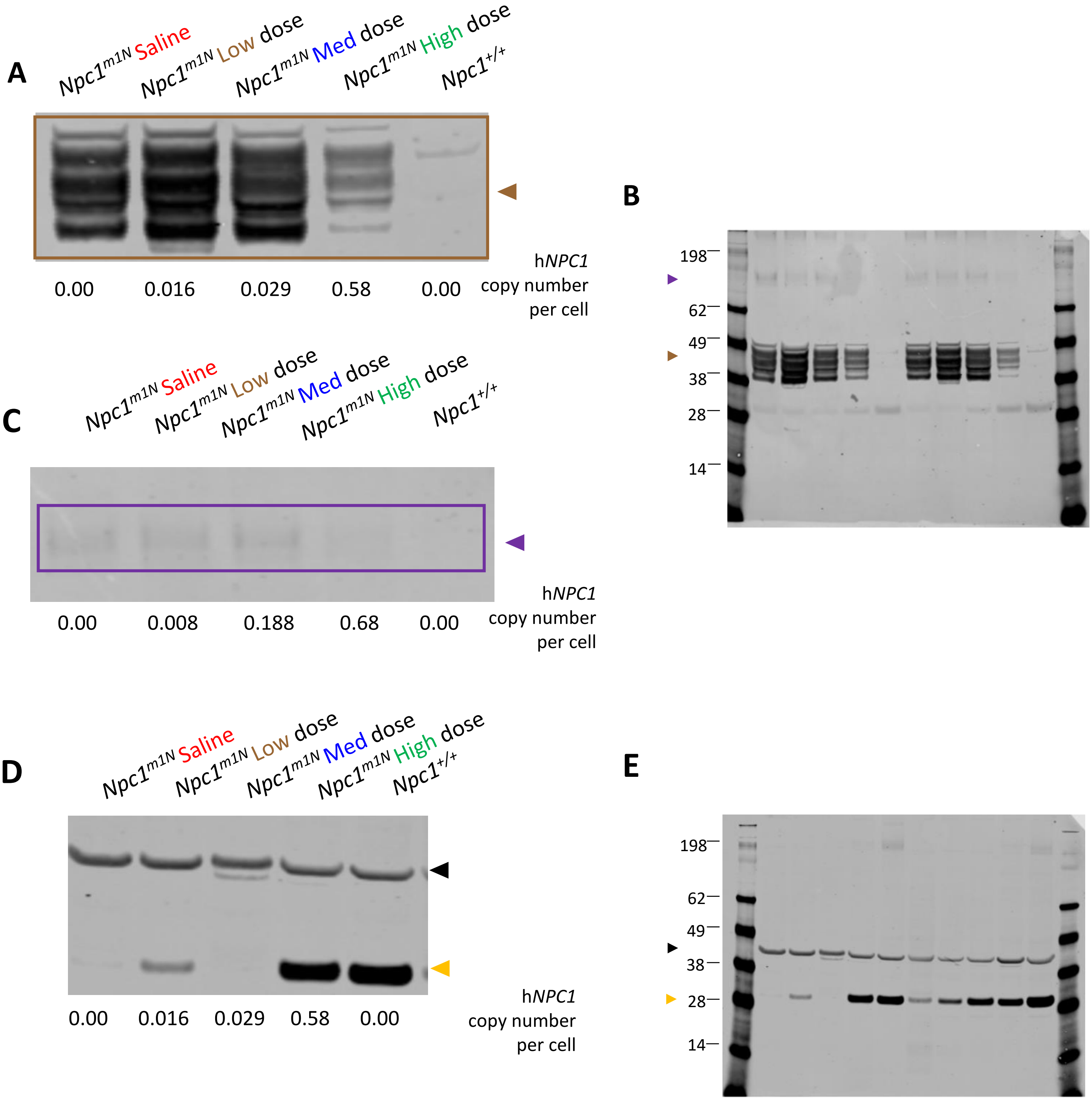
A
Npc1^m1N Saline
Npc1^m1N Low dose
Npc1^m1N Med dose
Npc1^m1N High dose
Npc1^+/+
0.00
0.016
0.029
0.58
0.00
hNPC1 copy number per cell
B
198
62
49
38
28
14
C
Npc1^m1N Saline
Npc1^m1N Low dose
Npc1^m1N Med dose
Npc1^m1N High dose
Npc1^+/+
0.00
0.008
0.188
0.68
0.00
hNPC1 copy number per cell
D
Npc1^m1N Saline
Npc1^m1N Low dose
Npc1^m1N Med dose
Npc1^m1N High dose
Npc1^+/+
0.00
0.016
0.029
0.58
0.00
hNPC1 copy number per cell
E
198
62
49
38
28
14

### Figure S5

A

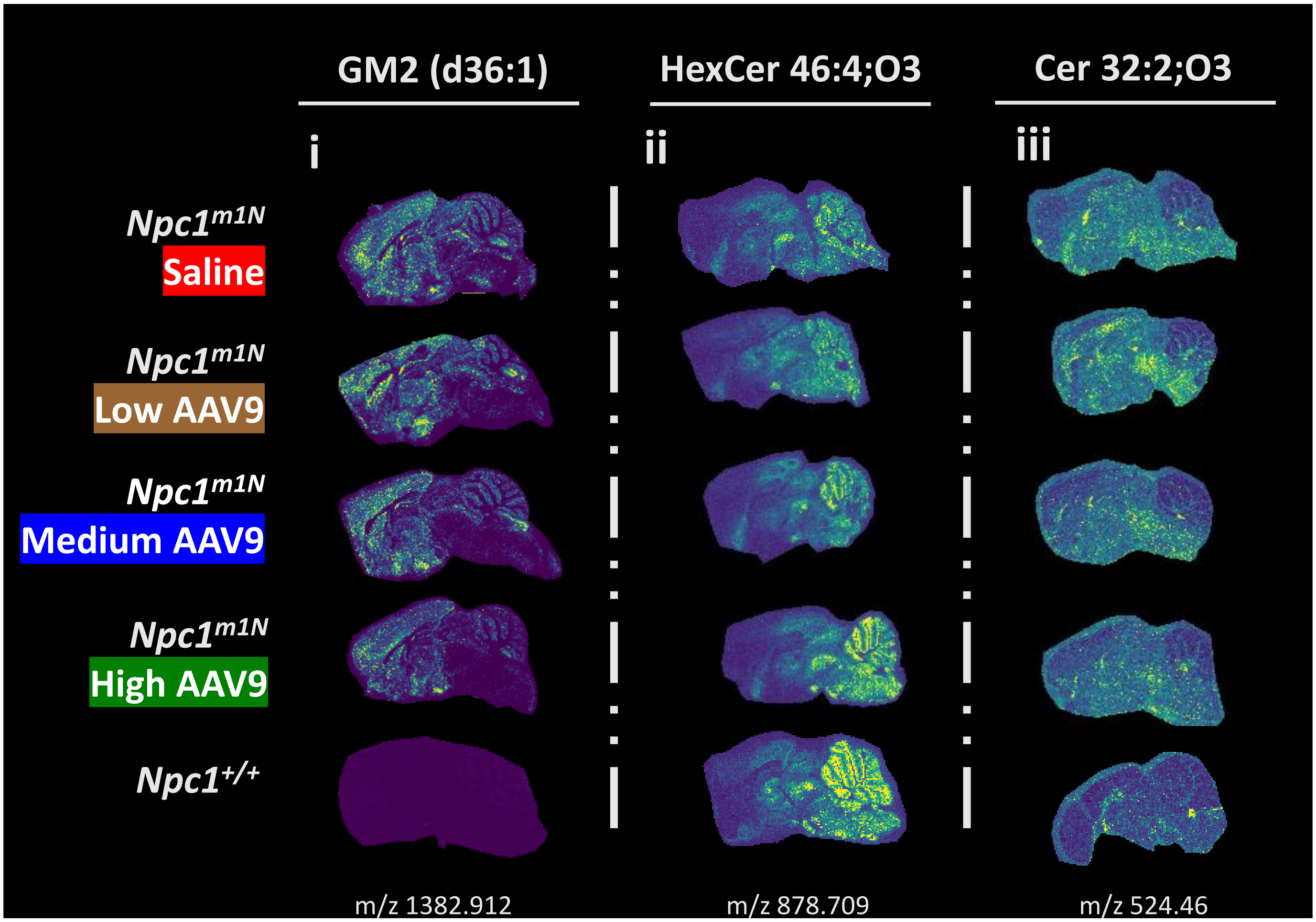

GM2 (d36:1)
HexCer 46:4;O3
Cer 32:2;O3
i
ii
iii
Npc1m1N Saline
Npc1m1N Low AAV9
Npc1m1N Medium AAV9
Npc1m1N High AAV9
Npc1+/+
m/z 1382.912
m/z 878.709
m/z 524.46


B

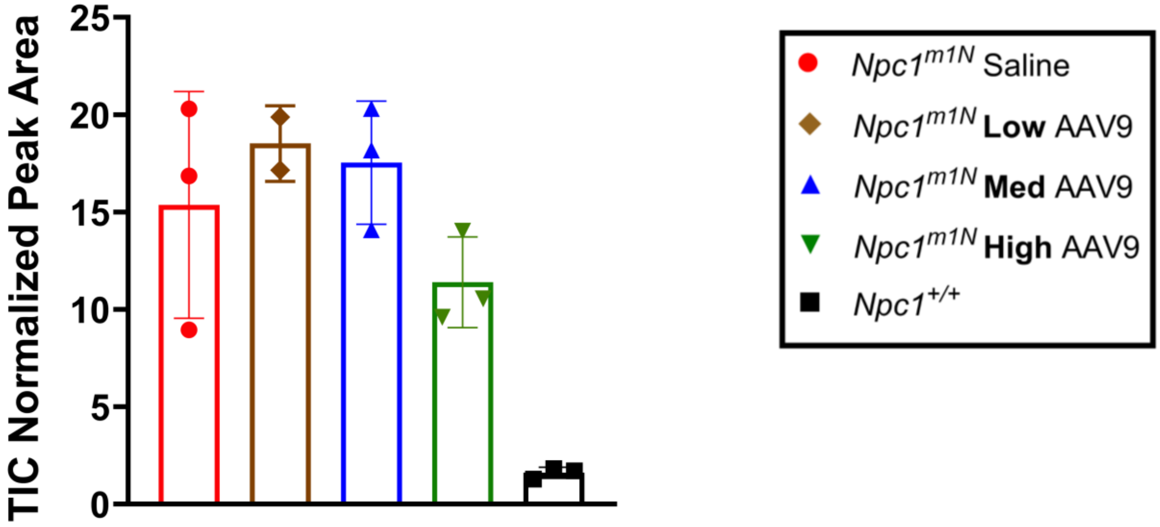

TIC Normalized Peak Area
25
20
15
10
5
0
Npc1m1N Saline
Npc1m1N Low AAV9
Npc1m1N Med AAV9
Npc1m1N High AAV9
Npc1+/+

**Figure S6**

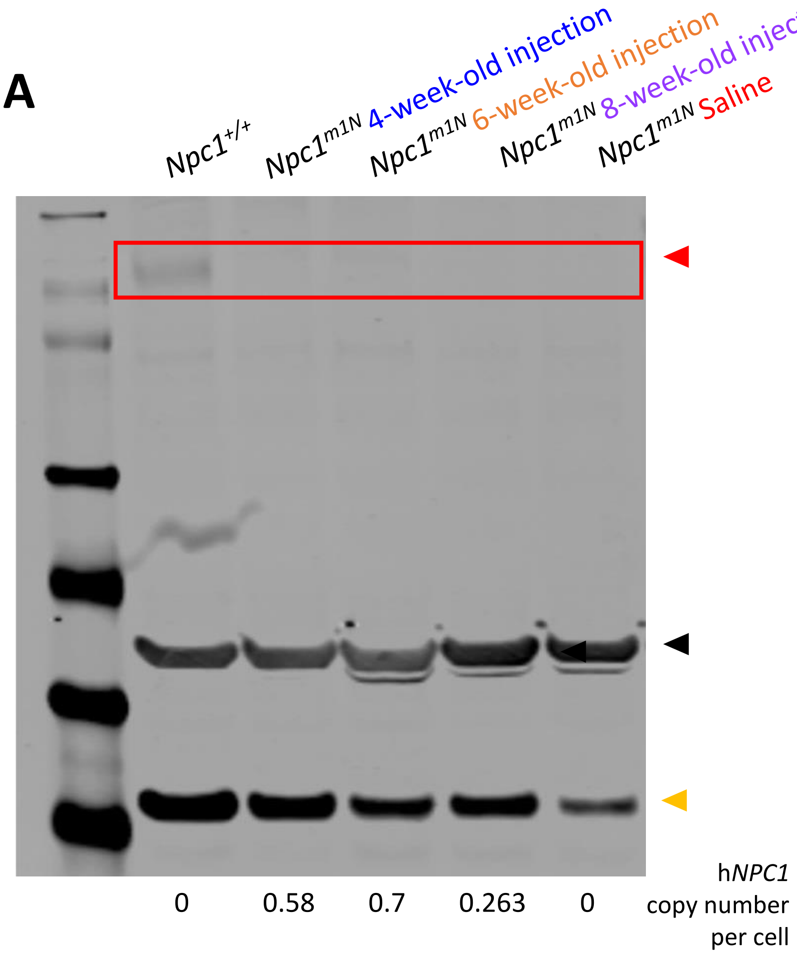
A
Npc1+/+
Npc1m1N 4-week-old injection
Npc1m1N 6-week-old injection
Npc1m1N 8-week-old injection
Npc1m1N Saline
0
0.58
0.7
0.263
0
hNPC1 copy number per cell

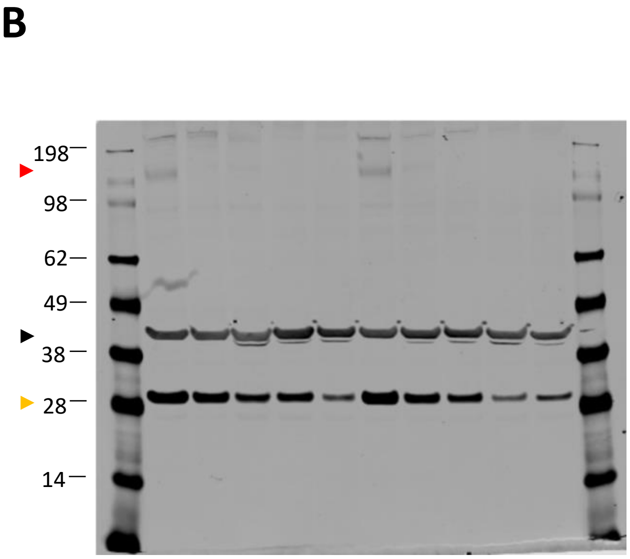
B
198
98
62
49
38
28
14

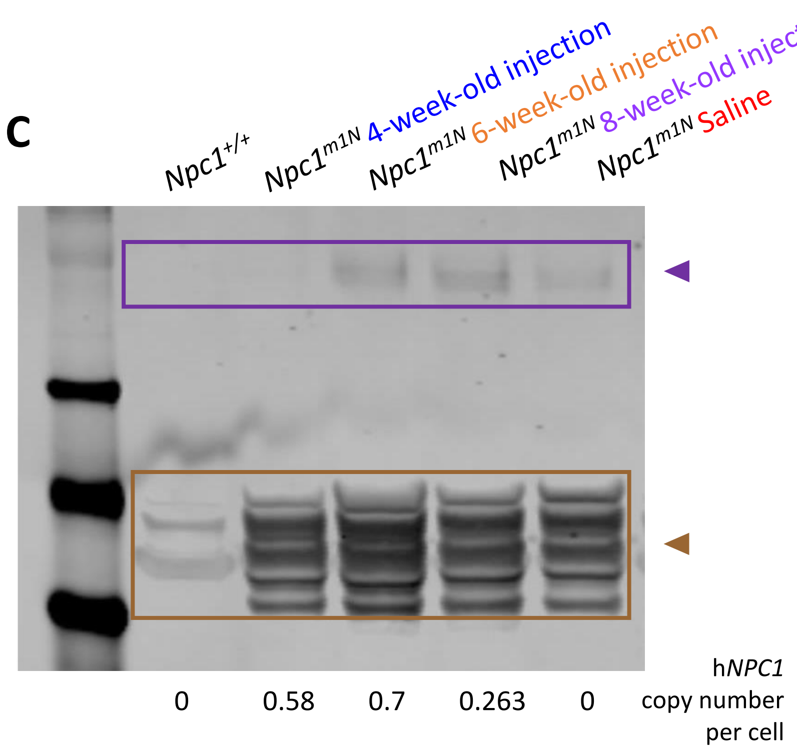
C
Npc1+/+
Npc1m1N 4-week-old injection
Npc1m1N 6-week-old injection
Npc1m1N 8-week-old injection
Npc1m1N Saline
0
0.58
0.7
0.263
0
hNPC1 copy number per cell

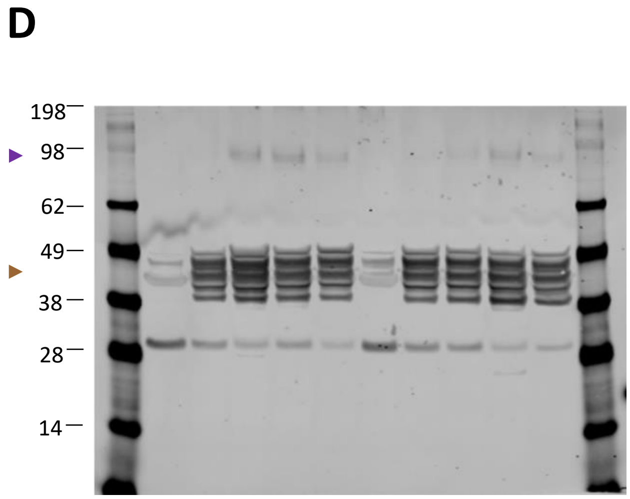
D
198
98
62
49
38
28
14

**Figure S7**

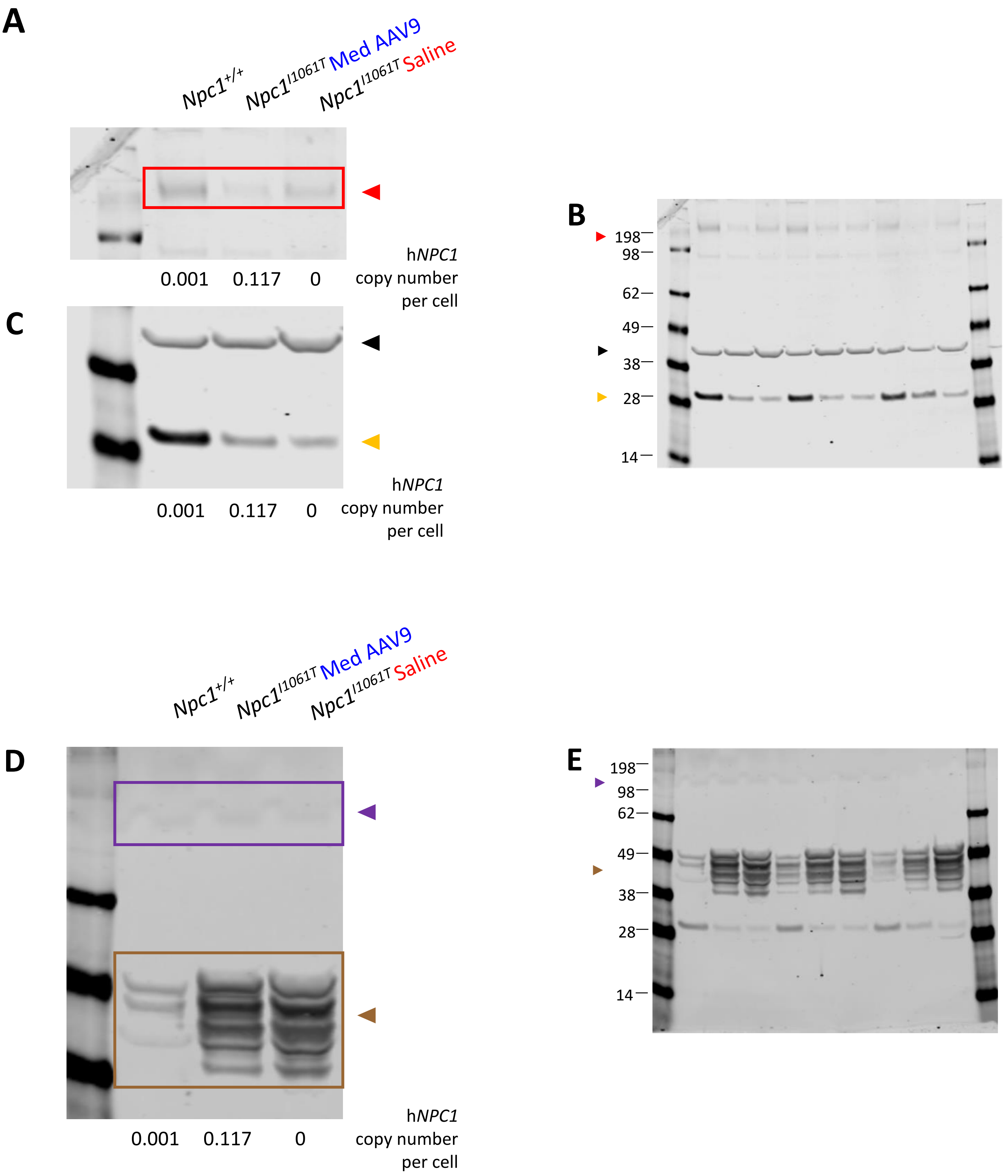

A
Npc1+/+
Npc1I1061T Med AAV9
Npc1I1061T Saline
0.001
0.117
0
hNPC1 copy number per cell
B
198
98
62
49
38
28
14
C
0.001
0.117
0
hNPC1 copy number per cell
Npc1+/+
Npc1I1061T Med AAV9
Npc1I1061T Saline
D
0.001
0.117
0
hNPC1 copy number per cell
E
198
98
62
49
38
28
14
